# Activation mechanism of Small Heat Shock Protein HSPB5 revealed by disease-associated mutants

**DOI:** 10.1101/2022.05.30.493970

**Authors:** Christopher N Woods, Maria K Janowska, Lindsey D Ulmer, Jasleen Kaur Sidhu, Natalie L Stone, Ellie I James, Miklos Guttman, Matthew F Bush, Rachel E Klevit

**Affiliations:** Department of Biochemistry, University of Washington, Seattle, United States; Department of Chemistry, University of Washington, Seattle, United States; Department of Medicinal Chemistry, University of Washington, Seattle, United States; Molecular Engineering and Sciences Institute, University of Washington, Seattle, United States

**Author notes:** Corresponding Author: Rachel E. Klevit. Equal contributions (Christopher N Woods, Maria K Janowska, Lindsey D Ulmer). Classification: Biological Sciences/Biochemistry.

## Abstract

Found from bacteria to humans, small heat shock proteins (sHSPs) are the least understood protein chaperones. HSPB5 (or αB-crystallin) is among the most widely expressed of the ten human sHSPs, including in muscle, brain, and eye lens where it is constitutively present at very high levels and carries out a myriad of functions. A high content of disorder in HSPB5 has stymied efforts to uncover how its structure gives rise to function. To uncover its mechanisms of action, we compared human HSPB5 and two disease-associated mutants, R120G and D109H. Expecting to learn how the mutations lead to loss of function, we found instead that the mutants are constitutively activated chaperones while wild-type HSPB5 can transition reversibly between non-activated (low activity) and activated (high activity) states in response to changing conditions. Techniques that provide information regarding interactions and accessibility of disordered regions revealed that the disordered N-terminal regions (NTR) that are required for chaperone activity exist in a complicated interaction network within HSPB5 oligomers and are sequestered from solvent in non-activated states. Either mutation or an activating pH change cause rearrangements in the network that expose parts of the NTR, making them more available to bind an aggregating client. While beneficial in the short-term, failure of the mutants to adopt a state with lower activity and lower NTR accessibility leads to increased co-aggregation propensity and, presumably, early cataract. The results support a model where chaperone activity and solubility are modulated through the quasi-ordered NTR and its multiple competing interactions.

**Significance:** Small heat shock proteins (sHSPs) are the oldest known protein chaperones, but how they recognize misfolding proteins in early stages of aggregation is unknown. Disordered regions within sHSPs are critical to their function, raising the question of how disorder recognizes disorder. We investigated a human sHSP (HSPB5) and two disease mutants using approaches that provide residue-level information in regions of disorder. The findings reveal that large, heterogeneous oligomers of HSPB5 contain complicated networks of interactions involving their disordered regions that protect them from solvent. Conditions associated with stress or disease mutations cause a rearrangement of the interaction network and enhancement of chaperone activity. The study provides new information regarding how disordered regions are modulated in a network of competing interactions.

## Introduction

Protein aggregation must be controlled under conditions of short-term acute stress and over long timeframes to maintain cellular health. Protein aggregates including fibrillar aggregates associated with neurodegeneration and amorphous aggregates associated with cataract are hallmarks of diseases that manifest in diverse tissues (Pedersen and Heegaard, 2013; Boatz et al., 2017). Small heat shock proteins (sHSPs) are ATP-independent chaperones that contribute to protein homeostasis by inhibiting protein aggregation (Horwitz, 1992). Although sHSPs were first discovered over 40 years ago (Ingolia and Craig, 1982), how the ten human sHSPs interact with diverse client proteins, their mechanisms of action, and how their activity is modulated remain poorly understood.

Named “small heat shock proteins” based on their polypeptide chain size (∼20 kDa), most sHSPs exist as large heterogenous and polydisperse oligomers containing ∼10-40 subunits (Baldwin et al., 2011a). Small HSPs share a common core domain, the α-crystallin domain (ACD), which is the only stably structured region within oligomers. The ACD forms an IgG-like β-sandwich structure that homodimerizes via a long β-strand to form the dimer interface (Bagnéris et al., 2009). Although the ACD dimer is thought to be the oligomeric building block, ACD dimers do not pack into discrete arrays or geometries in oligomers, giving rise to one level of oligomeric heterogeneity. All ACD dimer structures solved to date display two types of grooves on their surface, called “Edge” and “Central” for their positions on the dimer (Figure 1A). The ACD is flanked by disordered N- and C-terminal regions (NTR and CTR, respectively) that are variable in length and sequence and that contain small motifs that can bind weakly and transiently into ACD grooves (Figure 1A, Sluchanko et al., 2017; Clark et al., 2018; Clouser et al., 2019; Klevit, 2020). Although often required for the chaperone activity of sHSPs against aggregating clients, little is known about how disordered regions carry out the delay of aggregation, how they support differences in client preference, and how their activity is modulated under differing conditions.

**Figure 1.**
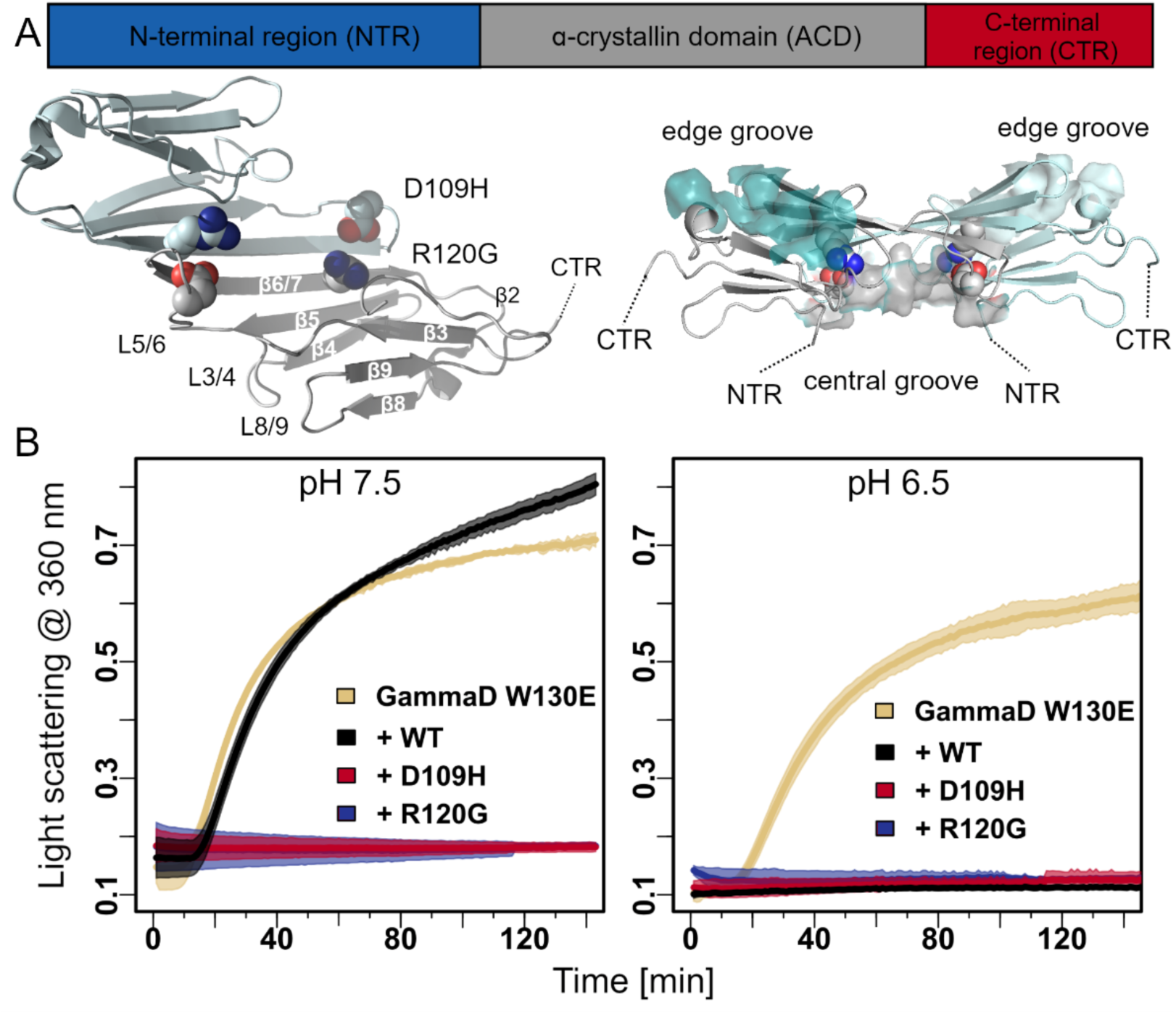
HSPB5 disease-associated mutants are activated chaperones against GammaD-crystallin W130E. A) **Top:** HSPB5 domain architecture includes an N-terminal region (NTR), α-crystallin domain (ACD), and C-terminal region (CTR). **Bottom:** Views of the ACD dimer (PDB 2N0K). *Left,* cartoon rendition of ACD dimer with disease mutation positions shown as spheres. One protomer of the dimer is labeled with its secondary structure elements. *Right*, Surface rendition with Central and Edge grooves highlighted by gray and cyan surfaces, respectively. Each dimer has two Edge grooves and one Central groove. B) GammaD aggregation assay. GammaD W130E (500 μM) was incubated alone (yellow) or in the presence of 100 μM WT HSPB5 (black), HSPB5 D109H (red), or HSPB5 R120G (blue) at 37 °C, pH 7.5 (*Left*) or at 37 °C, pH 6.5 (*Right)*. Aggregation was monitored by absorbance at 360 nm.

Mutations have been identified in human sHSPs that are associated with autosomal dominant inheritance of tissue-specific diseases that include neuropathy, cataract, and myopathy (Datskevich et al., 2012). The most extensively studied disease-associated mutation is substitution of Arginine 120 with Glycine (R120G) in ubiquitously expressed HSPB5, also known as αB-crystallin (Vicart et al., 1998; Fardeau et al., 1978). HSPB5 R120G is associated with desmin-related myopathy and early cataract (Vicart et al., 1998). The mutated residue is on the long β6+7-strand that associates with another copy of itself to form the ACD dimer interface and the Central groove (Figure 1A) (Bagnéris et al., 2009). Patient mutations of HSPB5 D109 (D109H, D109A, D109G) are associated with similar pathologies to R120G (Sacconi et al., 2012; Fichna et al., 2017; Brodehl et al., 2017). Intriguingly, D109 can form an ionic interaction with R120 in an adjacent ACD, suggesting a reciprocity of function (Figure 1A). Positions homologous to R120 and D109 are conserved in seven of the ten human sHSPs and are known sites of disease-related mutations, implying that what is learned for HSPB5 will be informative for a majority of human sHSPs (Datskevich et al., 2012).

Previous studies have characterized properties of HSPB5 R120G. Expression of HSPB5 R120G in cells or animals is associated with formation of aggregates and eye lens defects (Wu et al., 2018; Andley et al., 2011; Perng et al., 2004; Wang et al., 2001). Numerous studies have shown that HSPB5 R120G forms larger oligomeric assemblies that have more exposed hydrophobic surface than WT, faster subunit exchange dynamics, and are less effective at maintaining solubility of model client proteins under destabilizing conditions (Bova et al., 1999; Treweek et al., 2005; Liang and Liu, 2006). But the molecular details that underpin the deleterious consequences of the HSPB5 R120G mutation remain undefined.

The current picture of how deleterious mutations affect HSPB5 activity comes from studies using model client proteins. These tend to be either secreted proteins stabilized by disulfide bonds whose aggregation can be brought about by addition of reducing agents or proteins that are destabilized at temperatures above physiological (42 °C and higher). In such assays, R120G HSPB5 is a poor chaperone that often tends to co-aggregate with the client protein. It is unlikely that HSPB5 ever encounters these proteins or conditions. However, the tissue specificity of sHSP disease phenotypes implies that certain mutations are particularly deleterious under some conditions or toward certain intracellular client proteins. For example, mature lens fiber cells contain high concentrations of soluble, densely packed proteins known collectively as crystallins at concentrations that reach ∼400 mg/mL (Bloemendal et al., 2004). Differentiated lens cells lack the ability to synthesize or degrade proteins, so lens proteins accumulate damage over an individual’s lifetime (Lynnerup et al., 2008). Failure to prevent aggregation of damaged lens proteins, especially γ- and β-crystallins, results in cataract, a leading cause of blindness worldwide (Foster, 2000). Lens cells are also enriched in two sHSPs (HSPB4 and HSPB5; also known as αA- and αΒ-crystallin, respectively) that are present at an astounding subunit concentration of ∼7.5 millimolar. In light of the cataract-associated phenotypes of HSPB5 R120G and D109H, we focus here on a bona fide eye lens client protein, γD-crystallin (“GammaD”), which constitutes ∼10% of the protein mass in lens, for an approximate concentration of ∼2 mM (Robinson et al, 2006)

We compared WT and mutant HSPB5 using biochemical aggregation assays, hydrogen deuterium exchange coupled with mass spectrometry (HDX-MS), and photo-crosslinking coupled with mass spectrometry. Based on previous studies, we expected to learn how mutations in the structured ACD lead to a loss of chaperone function. Instead, we found that HSPB5 R120G and D109H exhibit constitutive activation towards a physiological client, GammaD. Furthermore, while WT HSPB5 can reversibly transition between non-activated and activated states in response to changing conditions, the mutant HSPB5s remain in an activated state. Our results reveal a complicated network of NTR-NTR and NTR-ACD interactions that changes upon chaperone activation. We offer a rationale for how a missense mutation in an ACD residue can bring about substantial rearrangement of the quasi-ordered NTRs, rendering them more exposed in the disease mutants and in activated WT HSPB5. Increased accessibility of certain NTR regions makes them more available to bind aggregation-prone clients but also increases the tendency of the small heat shock protein to aggregate on its own or co-aggregate with a client. We propose that long-term benefits of HSPB5 (and other sHSPs) requires an ability to transition reversibly between lower-activity, lower NTR-accessible states and higher-activity, higher NTR-accessible states. This transition can be driven by changes in the ACD dimer structure brought about by mutation (irreversible) or by a change in pH (reversible).

## Results

### Disease-associated mutants of HSPB5 are activated chaperones of a major eye lens protein client

The disordered NTR of HSPB5 is essential for its chaperone activity towards γD-crystallin (“GammaD”), one of the major protein clients in eye lens (Woods et al., 2023). In the absence of the NTR, the structured ACD displays no chaperone activity towards GammaD (Woods et al., 2023). In this study, we compared the activity of WT HSPB5 and two disease-associated mutations that target ACD residues to understand the interplay between the structured and disordered regions in chaperone function.

Like all crystallin proteins, GammaD is highly stable and does not normally aggregate at 37 °C, even at high concentrations. Therefore, we used GammaD W130E, a mutant that mimics oxidation-damaged GammaD that may build up over an individual’s lifetime. GammaD W130E is folded and can be purified but will spontaneously aggregate at 37 °C, both at pH 7.5 and pH 6.5 (Serebryany et al., 2016; Woods et al., 2023). These pH conditions span those observed in lens cells, depending on their placement in the lens and their age (Bassnett and Duncan, 1985; Mathias et al., 1991; Leem at al., 1999). On its own, GammaD W130E (500 μM) aggregates rapidly at both pH values, as evidenced by light scattering (monitored by absorbance at 360 nm). (Figure 1B, *left and right,* yellow trace). At pH 7.5 and at sub-stoichiometric levels (100 μM), WT HSPB5 had limited ability to delay the onset of aggregation, while HSPB5 R120G and D109H delayed aggregation for more than two hours (Figure 1B, *left,* black, blue, and red traces). In contrast, the three variants are all highly effective at delaying GammaD W130 aggregation at pH 6.5 (Figure 1B, *right*). Thus, WT HSPB5 adopts a non-activated conformation at pH 7.5 and an activated state at pH 6.5 but the mutants exist in an activated state regardless of pH conditions. To assess whether the mutants are better, worse, or equally effective under low pH conditions, aggregation assays were performed at a lower HSPB5 concentration (Supplemental Figure S1A). At low chaperone concentrations as may occur over an individual’s lifetime (Schmid et al., 2021), the disease mutants are less effective than WT HSPB5. Finally, aggregation assays performed in the presence of ACD-only and ACD-CTR constructs revealed that neither mutant ACDs nor the WT ACD (as previously reported) has chaperone activity towards GammaD W130E under either pH condition (Supplemental Figure S1B, C, D). Hence, the biochemical phenotype of the R120G and D109H mutations is only expressed in the context of intact oligomeric HSPB5 and a full understanding of the consequences of the two ACD mutations requires investigation of the full-length proteins as they present themselves in heterogeneous oligomers.

### Disease-associated mutations in ACD residues affect NTR structure and dynamics

To characterize differences in the biophysical properties of the disordered and highly hydrophobic NTR in the context of full-length oligomers, we performed hydrogen deuterium exchange coupled with mass spectrometry (HDX-MS). As shown previously, HDX-MS provides peptide-resolution information on the disordered NTRs of sHSPs within oligomers (Clouser, et al. 2019; Kaiser et al., 2019; Woods et al., 2023). HDX reflects the ability of protein backbone amide groups to exchange hydrogen with deuterium in the form of ^2^H2O. Dynamic or unstructured regions exchange rapidly (seconds-minutes), while structured, hydrogen-bonded, and/or buried sites exchange slowly (hours-days) and are described as “protected.” The degree of protection from exchange is indicative of differences in local structure and dynamics. We compared the HDX behavior of WT, R120G, and D109H HSPB5 at pH 7.5, conditions where the WT and mutants display the largest differences in activity, over a timescale of 4 sec to 20 hours (Figure 2). All datasets had 100% coverage across the sequences, although certain regions were represented only in fairly long peptides (Supplemental Table 1). Except in the ACD, where the proteolysis step (using immobilized pepsin and AN-PEP) produced long peptides, we focus on peptides of 12 residues or fewer, as these provide higher resolution information. Deuterium uptake plots for exemplary peptides that span the NTR and ACD are shown (WT, black; R120G, blue; D109H, red); in each case, similar peptides with slightly different ends showed similar behaviors (Supplemental Figure S2A). The CTR was only represented in long peptides that are completely exchanged after 4 seconds. (Data for all peptides are provided in Supplemental Figure S2A.)

**Figure 2.**
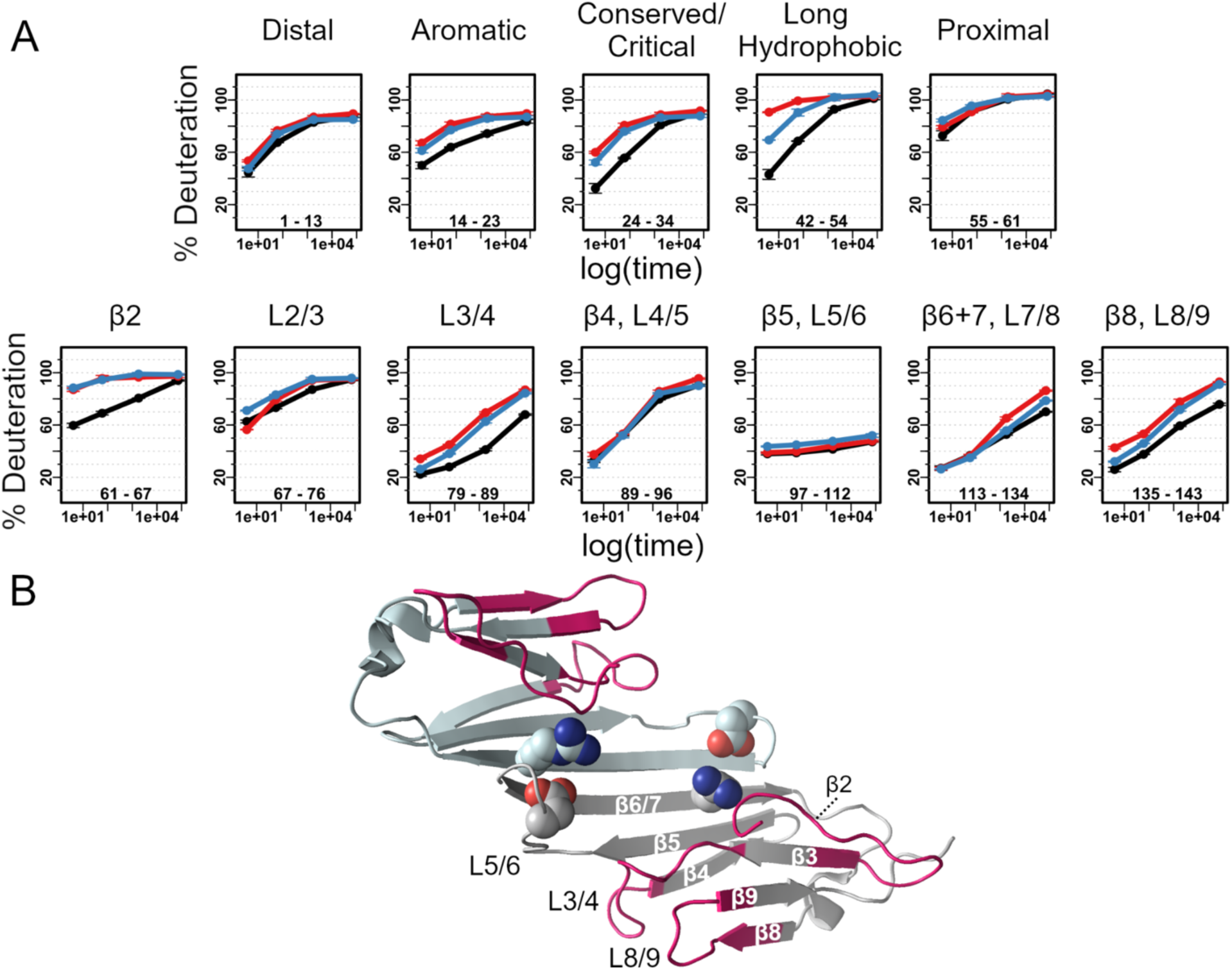
Hydrogen/Deuterium exchange reveals long-range effects of HSPB5 ACD mutations on oligomer structure and dynamics. A) Deuterium uptake plots for exemplary peptides that cover the HSPB5 sequence (WT, black; D109H, red; R120G, blue). Four time points were collected: 4 sec, 30 sec, 60 min, and 20 hr. The x-axis is plotted as time (sec) on a log scale. Top row, peptides that span the NTR are shown. Bottom row, peptides that span the structured ACD are shown. CTR-containing peptides are not shown, as they are completely exchanged at the first time point in all data sets. B) Structure of HSPB5 ACD dimer (PDB 2N0K), displaying the two mutated residues as spheres. Residues contained in the peptides that display increased deuterium uptake in the mutant oligomers are colored magenta.

As previously reported, the disordered NTR and structured ACD regions of HSPB5 display different deuterium exchange behavior. Although they lack defined structures, we can define six sub-regions within the NTR according to their positions relative to the ACD and their amino acid compositions: Distal (residues 1-13), Aromatic (residues 14-23), Conserved (residues 24-30), Critical (residues 31-35), Long Hydrophobic (residues 36-54), and Proximal (residues 55-61) (Figure 2A). The short Conserved and Critical regions have been previously identified: the Conserved region is highly conserved in sHSPs and the Critical region contains residues that when swapped with their respective amino acids from highly related HSPB4 is sufficient to switch HSPB5 chaperone activity to that of the less active HSPB4 (Woods et al., 2023).

Consistent with its H-bond-rich β-sheet structure, ACD peptides (Figure 2A, bottom row) are the most protected (lowest %D at 4 sec, the first timepoint) and some are incompletely exchanged after 20 hours (longest timepoint). Despite being disordered, most NTR-derived peptides in WT HSPB5 are less than 50% exchanged at 4 sec. NTR peptides (Figure 2A, top row) take up deuterium throughout the exchange time course and are fully deuterated by or before the 20-hour timepoint. No protection was observed for the disordered CTR, consistent with it being highly flexible and exposed. Altogether, the behaviors indicate that, although protected, the NTR regions undergo more frequent fluctuations to exchange-competent states than peptides derived from the structured ACD.

The mutated positions appear in different peptides in the HDX data: D109 is present in the β5, L5/6 peptide and R120 in the β6+7, L7/8 peptide (peptides are named for the secondary structure elements within the ACD; Figure 1A and 2B). Uptake for the peptide containing residue 109 (Peptide 97-112) is not perturbed in either mutant, while there is increased uptake for the peptide containing residue 120 (Peptide 113-134) at longer timepoints. The most affected ACD peptides have higher levels of deuterium uptake throughout the exchange time course (peptide β2, peptide L3/4, and peptide β8, L8/9; See Figure 2B). Different solved ACD structures have strand β2 patched to β3 in the top β-sheet of the β-sandwich structure or in a loop conformation; when it is patched, it sits atop R120 (Figure 2B). The peptide containing β2 residues is almost completely exchanged at 4 sec in both mutants, indicating that it is predominantly in a non-strand/loop conformation in mutant oligomers in solution. Loop L3/4 is proximal to R120 in the ACD structure and Loop L8/9 is adjacent to L3/4, suggesting a localized effect of mutating R120 on the secondary structure of these two loops or a change in their interactions with other regions of the oligomer. Intriguingly, these loops are slightly more affected in D109H HSPB5 although the Asp109 sidechain is not in direct contact with either affected loop in solved structures and is in fact in the neighboring subunit of the dimer (Figure 2C). This observation implies a linkage between the two positions, as mutation of either one affects other regions in similar ways.

Overall, NTR peptides of the mutants display substantially increased deuterium uptake compared to WT HSPB5 (Figure 2A, top row). The exception is the extreme N-terminal region (Distal; residues 1-13) where uptake plots of WT and mutants are very similar. The most highly protected peptides in WT HSPB5 are from the Conserved/Critical region (Peptide 24-37) and the Long Hydrophobic region (Peptide 42-54). These regions exhibit the largest changes in HDX in the mutants, with the Long Hydrophobic peptide almost completely deuterated at the 4 sec timepoint in D109H. The least protected peptides in the NTR contain residues that lead from the NTR into the ACD; these are almost completely exchanged at 4 sec in all three proteins. Thus, HDX reveals that the NTR of both mutant proteins is less protected, with the long Hydrophobic region being the most highly deprotected. The Distal peptide from the extreme N-terminal end remains sequestered and is the most protected NTR region in the mutant oligomers.

In summary, the two disease mutations located at the ACD dimer interface have similar effects on the dynamics of both the NTR and ACD regions of HSPB5 in the context of oligomers, with the effects of D109H being somewhat greater. In the ACD, the mutations appear to inhibit formation of the β2 strand that can patch onto the top β-sheet in WT HSPB5 to form part of the wall of the Central groove, and to affect two loops that are proximal to Arg120 and that lead into and out of one side of the Edge groove (Figure 2B). In the NTR, two of the most protected regions in WT HSPB5, namely the Conserved/Critical and the Long Hydrophobic regions are substantially de-protected in mutant oligomers, with the Long Hydrophobic region almost completely de-protected in D109H HSPB5.

### The NTR engages in a network of interactions

The HDX data at pH 7.5 point to dramatic rearrangements within the NTR of the mutant HSPB5 oligomers. The heterogeneity of HSPB5 oligomers defies conventional structural analysis, so we sought more detailed information using photo-crosslinking/mass spectrometry to map inter-chain interactions within oligomers. The non-canonical amino acid p-benzoyl-l-phenylalanine (BPA) was installed at single positions in HSPB5 using amber codon suppression (Chin et al., 2002). When excited at ∼360 nm, the benzophenone moiety of BPA forms covalent crosslinks to C-H bonds that are within 3.1 Å, providing a powerful probe for proximity/atomic contacts (Dorman and Prestwich, 1994).

We used a set of six BPA variants, in which BPA was substituted for an existing aromatic (or leucine in one case) residue in each of the NTR subregions: BPA-9 in Distal, BPA-17 in Aromatic, BPA-24 in Conserved, BPA-33 in Critical, BPA-47 in Long Hydrophobic, and BPA-61 in Proximal/β2 (Figure 3A) subregions. Irradiation was performed at pH 7.5 or at pH 6.5 for 30 minutes at 4 °C. As previously reported, SDS-PAGE analysis of the resulting samples revealed a major product at the expected position of two HSPB5 polypeptide chains attached via a single intermolecular cross-link (”dimeric products”).

**Figure 3.**
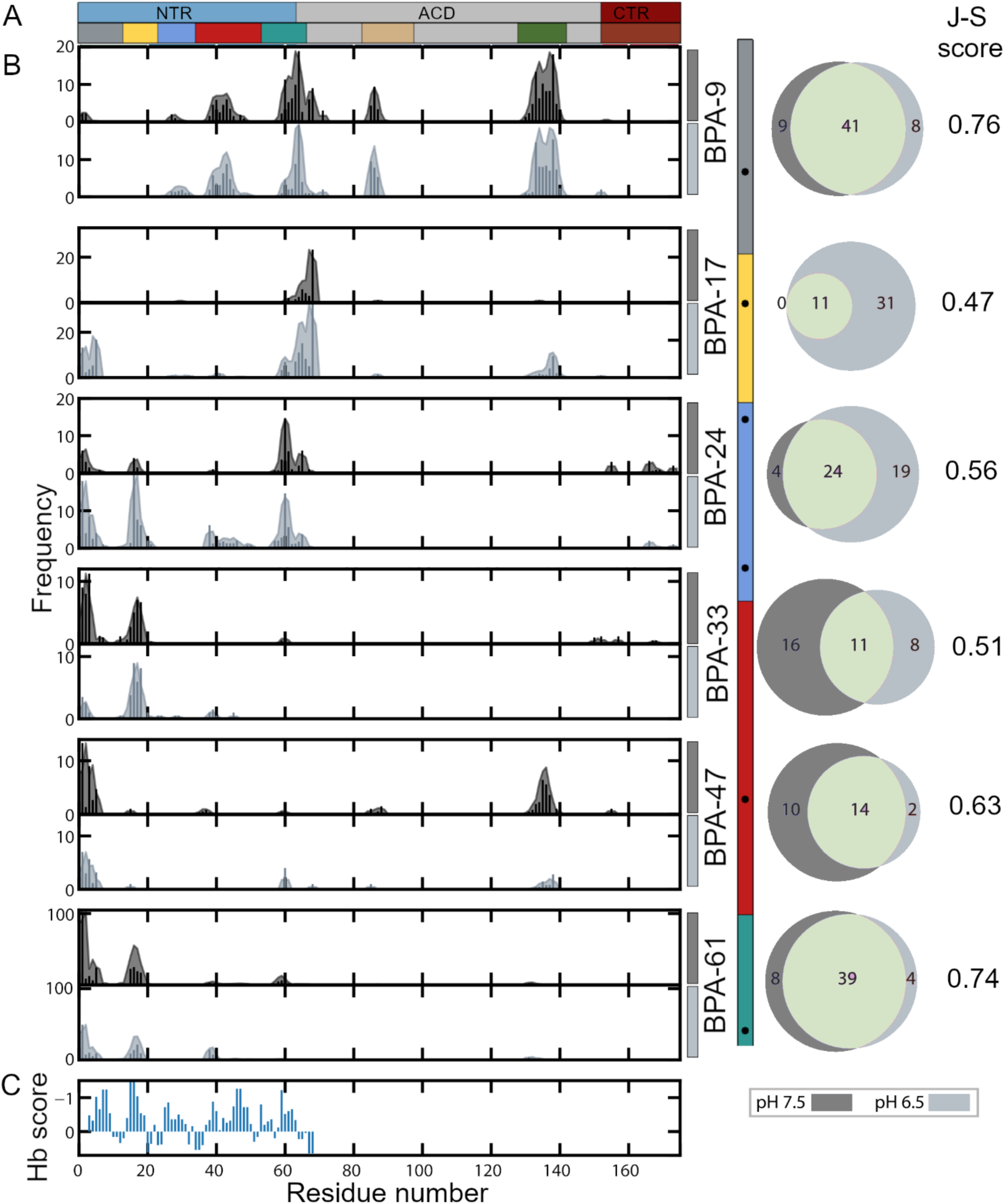
BPA cross-linking reveals HSPB5 NTR interactions. p-benzoyl-l-phenylalanine (BPA) was introduced into HSPB5 at residues 9, 24, 33, 47, or 61 (top to bottom). A) Positions of NTR sub-regions and β4- and β8 strands of the ACD are shown in colors. B) Histograms of cross-links between each BPA site and residues in HSPB5 identified at pH 7.5 (dark gray) and 6.5 (light gray) are shown. Bars indicate the total number of peptide spectral matches for each crosslink identified and shading shows the relative number of crosslinks in a 3-residue sliding window. The number of crosslink PSMs identified corresponds to how many spectra identified peptides with that crosslink site. The y-axis scale differs for each BPA site based on the number of crosslink PSMs identified (Supplemental Table 2 provides the total number of PSMs for each sample). Crosslinks for BPA sites 9, 17, 24, and 61 are from analyses of previously reported raw data (Woods, et. Al. 2023, *PNAS*) using updated versions of software (Ulmer, et. Al., 2024). To the right of each pair of histograms, a Venn diagram compares the crosslinks under the two pH conditions. The Jenn-Shannon similarity score for each BPA variant is given at the extreme right. C) Hydrophobicity according to Hopp & Woods, calculated with a 5-residue window is shown. More negative values are more hydrophobic (Note: y-axis is inverted for clarity).

Higher-order products representing >2 crosslinked subunits were also detected, but these were not investigated further (Woods et al., 2023 and Supplemental Figure 3A). Peptides generated by in-gel digests of the dimeric products were analyzed by mass spectrometry (Ulmer et al., 2024). The HSPB5 molecules must contain a covalent cross-link between two chains to be included in our analysis. Data-dependent acquisition and bioinformatic analysis of the resulting spectra identified multiple residues that were crosslinked. Results for each BPA variant are presented as histograms based on the observed peptide spectral matches (PSMs) for each identified crosslink (Figure 3B), with data for pH 7.5 and 6.5 shown in dark and light gray, respectively. We note that the number of PSMs should not be taken as a quantitative metric of the prevalence of a given interaction; we will interpret these qualitatively. To the right of each pair of histograms is the Venn diagram comparing the crosslinks identified under the two conditions. These will be discussed in the next section. Circle plots in which data from all variants are plotted, colored according to the sub-region or structural element in which the crosslink resides are shown in Figure 3C (or Supplementary Figure 3). Several points are immediately clear from the plots: 1) crosslinks are not randomly distributed, 2) contiguous stretches of residues are crosslinked with intervening sequences that gave no crosslinks, and 3) each BPA variant has a different pattern of crosslinks.

Focusing first on crosslinks from NTR sites to the structured ACD, some histograms have strong peaks around residue 138 (on β8 strand) and smaller peaks around residue 85 (on β4 strand). The Edge groove is created by strands β4 (residues 89-94) and β8 (residues 134-137) plus adjacent residues 85, 95, 98, 100, 130, 133, 143, and 145. Thus, BPA-9 (Distal) and BPA-47 (Long Hydrophobic) crosslink to the Edge groove. The Distal subregion contains the sequence Ile^3^-Ala-Ile^5^, a match to the “IxI” motif found in the CTR of many sHSPs that is known to bind to the Edge groove. There is no obvious IxI motif within the Long Hydrophobic region, but there are numerous methyl-containing residues that might bind into the two hydrophobic pockets in the groove. Thus, the photo-crosslinking/MS results identify two new Edge groove-interacting regions in the HSPB5 NTR in addition to the well-known CTR IxI motif. The only other ACD region identified in crosslinks contains residues 60-70, in which residues 66-69 can form β2, which is part of the Central groove at the ACD dimer interface (Figure 1A). Crosslinks to this region are observed for BPA-9 (Distal), BPA-17 (Aromatic), BPA-24 (Conserved), and BPA-61 (Proximal). The floor of the Central groove is created mainly by residues in the long β6+7 strands (residues 11-120). No crosslinks are observed to these residues, indicating that BPA at any of the positions used does not insert deep into the groove. Hence, among the six BPA HSPB5 derivatives, all but BPA-33 in the Critical region give crosslinks to the ACD and the Distal region contacts both grooves of the ACD.

Examination of NTR-NTR inter-subunit crosslinks reveals that each of the six BPA positions has a different crosslink pattern. Crosslinked residues are in contiguous patches along the NTR sequence and identify five interactive zones (Figure 3B): residues 1-6 (Distal), 12-18 (Aromatic), 25-32 (Conserved), 36-50 (Long Hydrophobic), and 57-63 (Proximal) (Figure 3A and B). In many cases, reciprocal crosslinks are observed: for example, BPA-17 crosslinks to residues 60-70 and BPA-61 crosslinks to residues 10-20, providing confidence that the incorporation of BPA has not disrupted native interactions. We note that the interactive zones identified by photo-crosslinking/MS correspond closely to five hydrophobic stretches according to a Hopp & Woods hydrophobicity analysis (Figure 3B, bottom panel). Although BPA reacts with any amino acid type, crosslinks were not identified in the short Critical region, which is an intervening hydrophilic stretch between the Conserved and Long Hydrophobic regions (Deseke et al., 1998).

The circle plot reveals a web of interactions captured in heterogeneous, polydisperse HSPB5 oligomers at pH 7.5 (Supplementary Figure 3, *left*). The picture that emerges is of multiple NTR regions that interact with the two known ACD grooves and a network of NTR-to-NTR interactions among the five hydrophobic zones. At pH 7.5, the Distal zone is the most highly interactive, contacting two ACD grooves and all NTR zones except the Aromatic. The Critical zone (BPA-33) crosslinks to the fewest number of regions, namely Distal and Aromatic. Altogether, the newly identified interactions involving the NTR greatly expand the limited information obtained from past solid-state NMR, cryo-EM, and biochemical studies (Ghosh and Clark, 2005; Jehle et al., 2010; Jehle et al., 2011; Peschek et al., 2013).

### HSPB5 Activation is accompanied by changes in its web of interaction

The HDX results presented in Figure 2 revealed that certain NTR regions are deprotected in R120G and D109H, implying that interactions that lead to their protection are lost or changed. Both mutants are activated chaperones at pH 7.5, so it is tempting to infer that the NTR, known to be required for chaperone activity, is released from sequestration in activated species (Figure 1B). To test this hypothesis, we repeated the photo-crosslinking experiments on all six BPA derivatives at pH 6.5, where WT HSPB5 is also activated (Rajagopal et al., 2015a; Woods et al., 2023). Identical workflows of sample irradiation, SDS-PAGE, in-gel digest, and MS analysis were carried out for the six BPA-HSPB5 derivatives at pH 6.5. Importantly, only the sample irradiation step was carried out under the two differing pH conditions; all other steps were carried out under identical conditions, so the resulting data sets can be compared across the two pH conditions. Histograms are shown in Figure 3B.

The total number of crosslink PSMs in an analysis depends on many factors, including the identity of the crosslinks in the sample, the quality of the experimental data, and the ability to identify the crosslinks in part due to the quality of the validation model. Therefore, the number of PSMs is not quantitative, and it is challenging to draw conclusions directly from the total number of crosslink PSMs for a sample. In contrast, the relative number of PSMs for a specific crosslinked product should be correlated with the relative abundance of the crosslink in the sample. Considering the relative abundance helps account for any differences in the total number of PSMs. However, if there is a large difference in the number of PSMs detected, it may still be difficult to compare because it becomes more likely that not all interactions are detected with fewer total PSMs. Here, we use the relative proportion of crosslinks to different regions to compare changes in the distribution of crosslinks for each BPA variant separately at two pH values to determine how NTR sub-region changes with pH.

We compared datasets collected at the two pH conditions in several ways. A Jenson-Shannon Divergence analysis was performed for each BPA variant (Lin 1991; Menendez et al., 1997). The analysis measures the similarity between the probability distributions associated with different samples (i.e. the normalized histograms of identified crosslinks) and provides a score between 0.0 and 1.0, where a higher score corresponds to higher similarity between the two samples. BPA-9 (Distal) and BPA-61 (Proximal/β2) have the highest scores, indicating that their crosslink patterns differ the least between pH 7.5 and pH 6.5. BPA-17 (Aromatic) has the lowest score (below 0.5), signifying that its pattern differs the most between non-activated and activated states. The other three positions show intermediate scores. Venn diagrams of the crosslink PSMs (Figure 3D) provide another way to compare the interactions of each BPA site in the two pH conditions. Consistent with their high similarity scores, the diagrams for BPA-9 and BPA-61 show a majority of each set overlapping with a small number of unique crosslinks at each pH. In contrast, the BPA-17 Venn diagram shows a large expansion of crosslinks in the activated state while the diagrams for BPA-33 and BPA-47 show a loss of crosslinks in the activated state.

We also sought finer-grained information from the data by comparing the sub-regions and structural elements involved in crosslinks for each BPA variant under the two conditions (Supplemental Table 2 and Figure 4). In both states, ∼95% of the crosslinks observed from BPA-9 (Distal) are to three regions, consistent with its high similarity score. In all other cases, both the number of regions involved in crosslinks and the identity of those regions differed between the two pH conditions. The most dramatic example is for BPA-17 (Aromatic), with >90% of its crosslinks going to a single region at pH 7.5 while they are distributed among three regions at pH 6.5. BPA-17 has the largest difference in the total PSMs between the pH values, with many more at pH 6.5 (45 PSMs versus 169 PSMs). While having fewer PSMs could result in detecting crosslinks to fewer regions due to the low number of observations, the opposite effect was observed for BPA sites 33, 47, and 61. In those samples, fewer crosslink PSMs were detected at pH 6.5 but these involved more zones of interaction, suggesting that detecting fewer PSMs is not significantly affecting our ability to detect a wide variety of crosslinks. Overall, the comparison reveals that there is substantial rearrangement of the complicated interaction network in activated versus non-activated HSPB5. Although the details of such rearrangements will require further investigation, one important point is clear: regardless of whether the total numbers of PSMs increased or decreased under activating conditions, the number of zones with which each BPA variant except BPA-9 interacts is higher. We rationalize this observation as follows. The interactions captured in BPA crosslinks identify the zones amongst which a given BPA residue is distributed. NTR interactions are weak and are essentially in competition with each other. A change in the number of competing interactions under differing conditions therefore suggests that the relative affinities of the relevant interactions have changed, allowing the BPA residue to partition and exchange among a larger number of zones. This interpretation is consistent with the observation of increased deuterium uptake of certain NTR regions in the constitutively activated mutant forms of HSPB5, as interchange between sites of interaction may occur via exchange-competent states, yielding de-protection against deuterium exchange.

**Figure 4.**
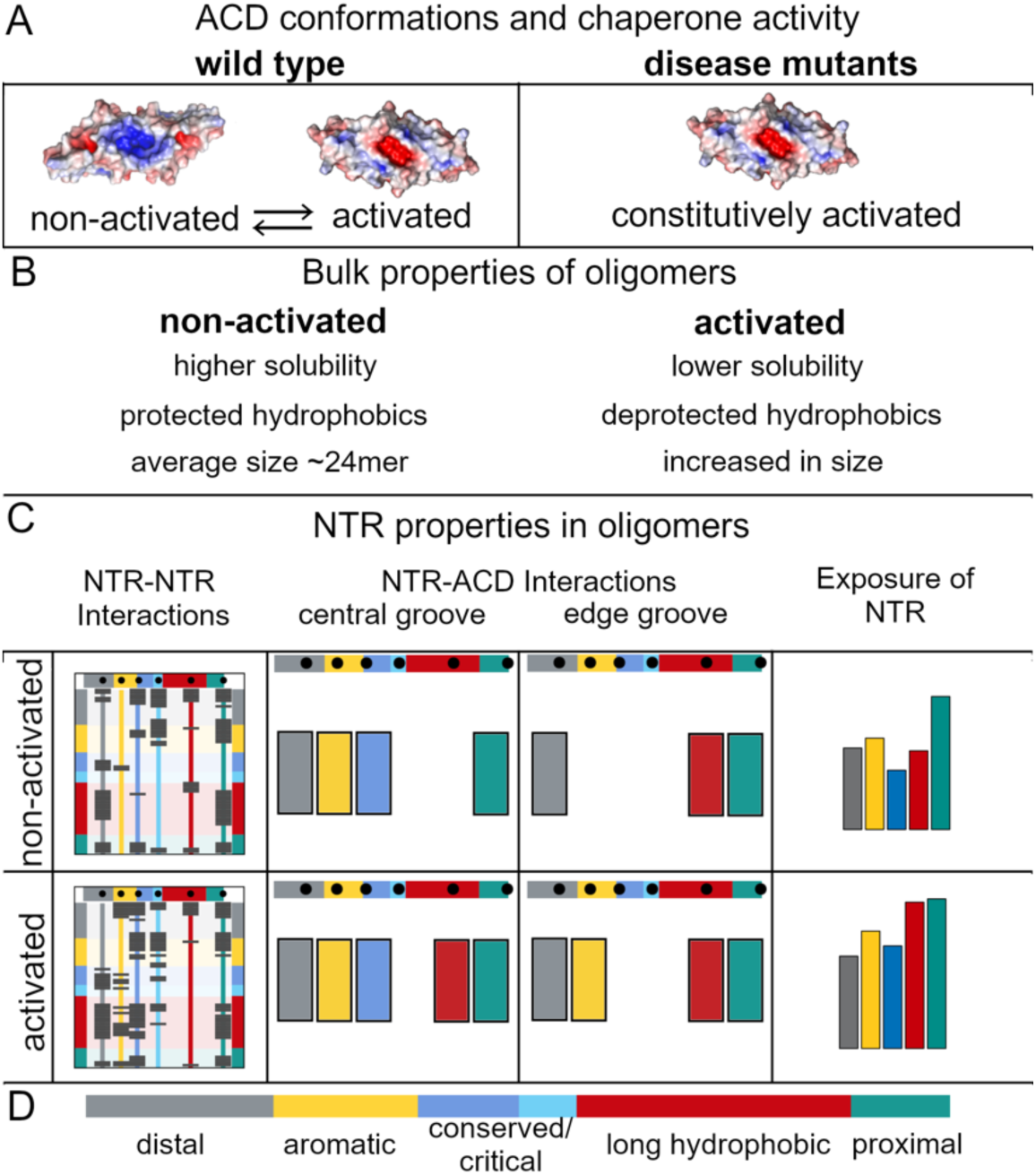
Features of the Activated State of HSPB5. A) ACD dimer structures differ between non-activated and activated HSPB5. (*Left*) WT transition reversibly between a non-activated dimer conformation (PDB 2N0K) and an activated conformation (PDB 2Y1Z). (*Right*) Disease mutants D109H and R120G ACD dimers adopt the activated conformation. Electrostatic potentials are shown on surface renditions, highlighting the large and positive-potential Central groove of the non-activated conformation versus the smaller and negative-potential groove of the activated dimer conformation. B) Comparison of the bulk properties of oligomers in non-activated and activated state. C) NTR properties within oligomers. Schematic presentation of the interactions (left and middle panels) and degree of exposure (right panel). Dark gray boxes on each colored vertical line in the “NTR-NTR Interactions” panels denote interactions identified from BPA crosslinking. Compare, for example, the yellow vertical line (interactions from the Aromatic region) in non-activated and activated states: there are many new interactions observed in the activated state. Colored bars in the panels labeled “NTR-ACD interactions” identify NTR regions that interact with each of the two ACD dimer grooves in activated and non-activated oligomers. D) Coloring used for the NTR sub-regions.

## Discussion

Small heat shock proteins are poorly understood compared to other chaperone proteins due to their functional promiscuity and structural heterogeneity. Despite its ubiquitous expression, some inherited missense mutations in sHSP HSPB5 are linked specifically to early cataract and certain myopathies (Datskevich et al. 2012). Many studies have focused on how one such mutation, HSPB5 R120G, affects the chaperone. These studies revealed major macroscopic and functional differences such as larger size, more exposed hydrophobics (Bova et al, 1999; Kumar et al., 1999; Perng et al., 1999), different shapes when viewed by EM, and formation of cytoplasmic and perinuclear aggregates when expressed in mammalian cells (Ito, et al. 2003) (Figure 4B). Thus, the structure of HSPB5 R120G is clearly perturbed but details on how it is changed have remained to be defined. A less-studied mutant, D109H HSPB5, is associated with the same narrow panel of tissue-specific diseases, raising the specter of a shared mechanism of action.

Previous studies found that HSPB5 R120G has decreased chaperone capacity or even enhances aggregation of model clients (Bova et al., 1999; Kumar et al., 1999; Perng et al., 1999). In contrast, we found both mutants have enhanced chaperone activity towards an important lens client, GammaD-crystallin, under physiological conditions. This apparent conflict may be due in part to the fact that HSBP5 R120G is less stable and less soluble at temperatures above 37 °C, where many model client aggregation studies are performed. At 37 °C, we found that WT HSPB5 has low chaperone activity pH 7.5 and increased activity at lower pH conditions that exist in some lens cells. The mutants are highly active under both conditions (Figure 1B). Thus, WT HSPB5 can transition between non-activated and activated states over a narrow pH range, presumably with intermediate levels of activity at pH values between the two extremes studied here. In contrast, the disease mutants are constitutively active towards GammaD (Figure 4).

That the disease mutants are permanently shifted towards a more activated state of HSPB5 raises the question, Are the activated states adopted by WT HSPB5 at low pH and the mutant proteins structurally similar? Slingsby and colleagues have noted that the structure of an R120G ACD dimer solved at pH 7.5 shares features with WT HSPB5 ACD dimer solved at low pH (Clark et al., 2011). The dimer interface in both is altered similarly, yielding a Central groove that differs from that in WT ACD at pH 7.5 in its size, shape, and electrostatics. Specifically, the groove is smaller, more closed, and has a negative electrostatic potential (Figure 4). No structures have been determined for HSPB5 D109H ACD dimer, but the HDX behavior reported here closely mirrors that of R120G HSPB5, consistent with similar structural perturbations. Although ACD dimers lack detectable chaperone activity towards GammaD, our results show that their structures have widespread and profound effects on the properties of the disordered NTRs that dictate chaperone activity. Therefore, to fully understand how a mutation within the ACD drives the overall conformational states of oligomers, it is incumbent to define how plasticity at the ACD dimer interface affects the properties and disposition of the disordered NTRs.

The high level of heterogeneity among and within HSPB5 oligomers defies their full resolution. We used two approaches to obtain residue- or peptide-level information on disordered NTRs as they exist in oligomers (Figure 4). Photo-crosslinking/MS identified six highly interactive regions with the NTR, punctuated by short hydrophilic, non-interactive sequences (Figure 3). HDX/MS revealed a general deprotection of NTRs in activated oligomers, except for the extreme N-terminal Distal region which remains protected. At the other end of the NTR, the Long Hydrophobic and Proximal regions are almost completely deprotected (Figures 2 and 4). Comparison of the photo-crosslinking in non-activated and activated species revealed substantial rearrangements in all regions of the NTR except the Distal region.

The two regions that differ the most between the states in their interaction network are the Conserved and Critical regions, in the middle of the NTR. The degree to which the Critical region in HSPB5 is protected/deprotected has been linked to chaperone activity in HSPB5 and its closest homolog, HSPB4 (Woods et al., 2023). Taken together, the results are consistent with a reorganization of the quasi-ordered interaction network in HSPB5 oligomers that is brought about by either a change in pH or mutation of residues at the ACD dimer interface.

Findings reported here are relevant to other members of the sHSP family in humans. Structures of ACD dimers from HSPB1, B2, B3, B4, B5, and B6 are highly similar and contain the two surface grooves, the Central and Edge grooves, discussed here. The first ∼30 residues of the NTR are resolved in a structure of full-length HSPB6, revealing its Distal region inserted into an ACD Edge groove and its conserved region inserted into the Central groove (Sluchanko et al., 2016). Most sHSPs, including those listed here, contain the Conserved region in their NTRs, implying similar Conserved-to-Central groove interactions.

Although there is no other discernable conservation within the NTR sequences, the pattern of hydrophobic/hydrophilic regions is conserved, suggestive of a shared architecture of interactive zones (Figure S4) and, hence, of interactive networks. For example, HSPB1, HSPB4, HSPB5, and HSPB6 all contain “IxI-like” motifs within their Distal regions that can interact with ACD Edge grooves. Inspection of other sHSP sequences indicates this is another shared feature and may serve to limit the range or reach of the highly hydrophobic NTRs under conditions that favor release.

In sum, we propose a model in which the well-known interactions between two ACDs generate dimers that are the building block of sHSP oligomers modulate quasi-ordered interactions with NTRs. Plasticity and dynamics at the dimer interface due to mutations and/or changes in pH lead to reorganization of the NTR interaction network and, consequently, modulation of chaperone activity. We imagine that mutations or post-translational modifications such as phosphorylation of NTR residues also lead to reorganization of the network, but details of these effects must await future study. sHSPs have evolved to undergo reversible transition between higher and lower activity states, but the mutations associated with disease in HSPB5 studied here lead to constitutive activation that is detrimental over the course of an organism’s lifetime.

## Materials and Methods

### Expression and purification of HSPB5 WT and mutants

*E. coli* BL21 cells were transformed with HSPB5 WT or mutants in a pET23 vector and expressed and purified as previously described (Woods et al. 2023). The same procedures were used for the HSPB5 ACD-CTR and ACD-only constructs, except a 120 mL SDX75 column was used for the final step. HSPB5 BPA-expressing cells were cultured identically to HSPB5 WT following induction by 1 mM IPTG and 1 mM Arabinose and purified as previously described (Woods et al., 2023).

### Expression and purification of GammaD-crystallin

GammaD-crystallin was expressed in *E. coli* BL21 cells as a C-terminal HexaHis-Sumo fusion in pET28a vector and purified as previously described (Woods et al., 2023). Fractions containing GammaD were concentrated and stored in 25 mM NaPO4, 150 mM NaCl, 0.1 mM EDTA pH 6.5 or 7.5.

### Chaperone Activity Assay

Chaperone activity assays were conducted in a 96-well flat-bottom half-area plate with a final volume of 100 μL. GammaD-crystallin was adjusted to a concentration of 1 mM and incubated on ice. Assay wells containing HSPB5 in 25 mM NaPO4, 150 mM NaCl (or buffer only) in 50 μL total volume were pre-heated for 5 minutes at 37 °C. Following pre-heating, 50 μL ice cold GammaD W130E in 25 mM NaPO4, 150 mM NaCl, 0.1 mM EDTA was added. The plate was incubated at 37 °C and A360 measurements were taken once per minute, with 5 seconds of shaking before each measurement.

### Hydrogen/Deuterium Exchange Mass Spectrometry

All samples were incubated at 37 °C for 3 hours in water-based buffer and then cooled to room temperature before undergoing deuterium exchange. HSPB5 WT and mutants (0.04 mg/mL) were incubated at room temperature in 85% D2O-based PBS pH 7.5 buffer for 4 seconds, 1 minute, 30 minutes, or 20 hours. Fully deuterated samples were made by incubating protein in 85% D2O buffer at 75 °C for 30 mins. Exchange was quenched by addition of ice-cold quench buffer (1.6% formic acid) and samples were flash frozen in liquid nitrogen. Samples were automatically thawed, digested by immobilized pepsin and AN-PEP (Tsitsiani et al., 2017), and injected on a Waters Synapt G2-Si instrument using a setup built in house around the LEAP PAL system (Watson et al., 2021). HSPB5 peptides were identified by MS/MS on a Thermo Orbitrap Fusion Tribrid instrument and MS^E^ on a Waters Synapt G2-Si followed by data analysis using ProteinProspector (UCSF) or ProteinLynx Global SERVER (Waters). Deuterium uptake was analyzed in HDExaminer 3.0 (Sierra Analytics) and HX-Express (Guttman et al., 2013). Multiple datasets were compared and statistically analyzed using HDXBoxeR (Janowska et al., 2024).

### BPA Crosslinking Mass Spectrometry

The BPA crosslinking experiments were performed as described previously (Woods et al., 2023 and Ulmer et al., 2024). The crosslinks for BPA sites 9, 17, 24, and 61 at pH 7.5 are analyses of previously reported raw data (Woods, et. Al. 2023, *PNAS*) using updated versions of software. The analyses of all datasets using the same software versions, with the exception of BPA at site 33 and pH 6.5 have been previously reported (Ulmer, et. Al., 2024, *JPR*). Data were collected with an Easy Nano LC coupled to a Thermo Orbitrap Fusion Lumos Tribrid Data was collected using data-dependent acquisitions and an 85-minute gradient using the previously reported mass spectrometry method. (JPR paper) The mass spectrometry proteomics data have been deposited to the ProteomeXchange Consortium via the PRIDE partner repository (Perez et al., 2019) with the dataset identifier PXD055848.

### Identification of Crosslinks Using the Trans Proteomic Pipeline (TPP)

Crosslinks were identified using the previously reported informatics workflow (Ulmer et al., 2024) along with the Trans Proteomic Pipeline version 6.3.2 (Deutsch, et al., 2023). Briefly, the method uses Comet (Eng et al. 2013) to search for non-crosslinked peptides in the monomeric reactant sample and uses PeptideProphet (Keller et al., 2002) to validate to a 1% false discovery rate (FDR) at the PSM level. The proteins identified are then used to create the validated protein database, which is used as the protein database to analyze the dimeric product sample. Dimeric product samples are searched using Kojak (Hoopmann et al., 2015) and the validated protein database. PeptideProphet is used to validate results to a 1% FDR at the PSM level. For histograms, each PSM was associated with the residue that was assigned the highest probability of participating in a crosslink with BPA. When more than one residue was assigned the same probability, an equal fraction of that PSM was assigned to each of those residues. The code used to assign residue-level crosslinks and create histograms is publicly available (https://github.com/bushgroup/Identifying-Site-Specific-Crosslinks) and was described previously. (Ulmer et al., 2024). Jenson-Shannon scores were calculated using the relative proportion (percent of total) crosslink PSMs at each residue in HSPB5.

## Acknowledgements

We thank Edgar A. Hodge, Mark A. Benhaim, and Clint Vorauer for assistance with HDX feasibility studies and protease immobilization. We thank Daniele Canzani for assistance with early BPA cross-linking mass spectrometry experiments and Ponni Rajagopal for foundational studies on R120G-HSPB5 ACD. We thank Peter Brzovic and Lisa Tuttle for helpful discussions and careful reading of the manuscript. This work was supported by NIH grants R01 EY017370 to REK and R01 AI153191 to MG, and by T32 GM008268 to CNW, T32 GM007750 to EIJ, T32 AG066574 to MKJ, and T32 AG066574 to LDU and by the University of Washington’s Proteomics Resource (UWPR95794).

## Competing Interests

The authors declare no competing interests.

## Notes

### Competing Interest Statement

The authors have declared no competing interest.

### Summary of Updates

In this revision, we compare properties of HSPB5 activated by disease mutations with WT-HSPB5 activated by pH. To enhance rigor and confidence, we have optimized analysis and bioinformatic workflows for the HDX/MS data and photo-crosslinking/MS data. The updated analysis reveals a shared mechanism of activation in which certain regions of the intrinsically-disordered NTR are released upon activation.

