## Supplemental Figures and Table for "Activation mechanism of Small Heat Shock Protein HSPB5 revealed by disease-associated mutants"

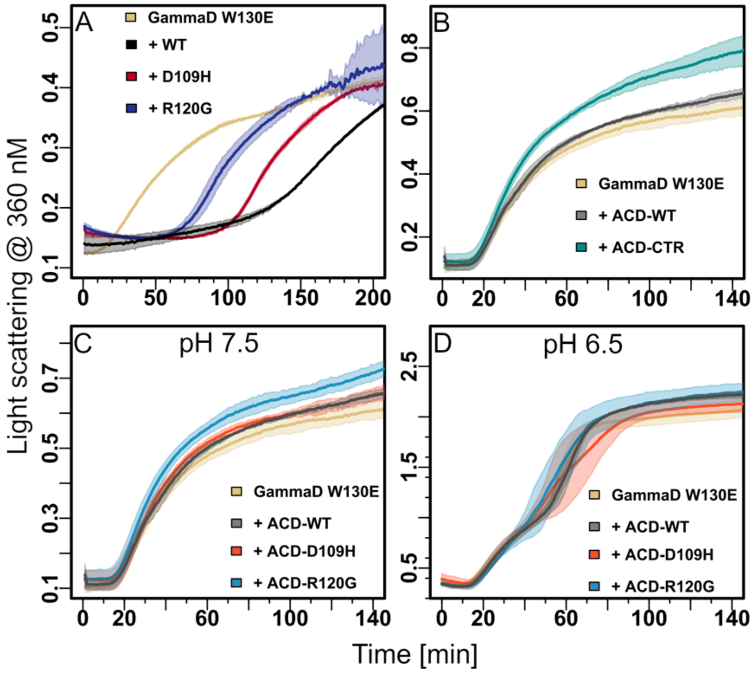


Fig. S1. Aggregation assays of GammaD. A) At pH 6.5, where WT- and both mutants are activated (Figure 1B), R120G- and D109H-are less effective chaperones when present at low concentration. Aggregation assays at pH 6.5 were conducted with 500uM GammaD and 2 uM sHSP. B, C, D) ACD dimers do not delay GammaD aggregation. Panel B compares the activity of WT-ACD dimers and WT-ACD-CTR dimers. Panel C and D compare WT-, R120G-, and D109H-ACD dimers at pH 7.5 and pH 6.5. All assays include 500 µM GammaD except in Panel D where 300 µM GammaD was used. All ACD constructs were present at 100 µM GammaD.

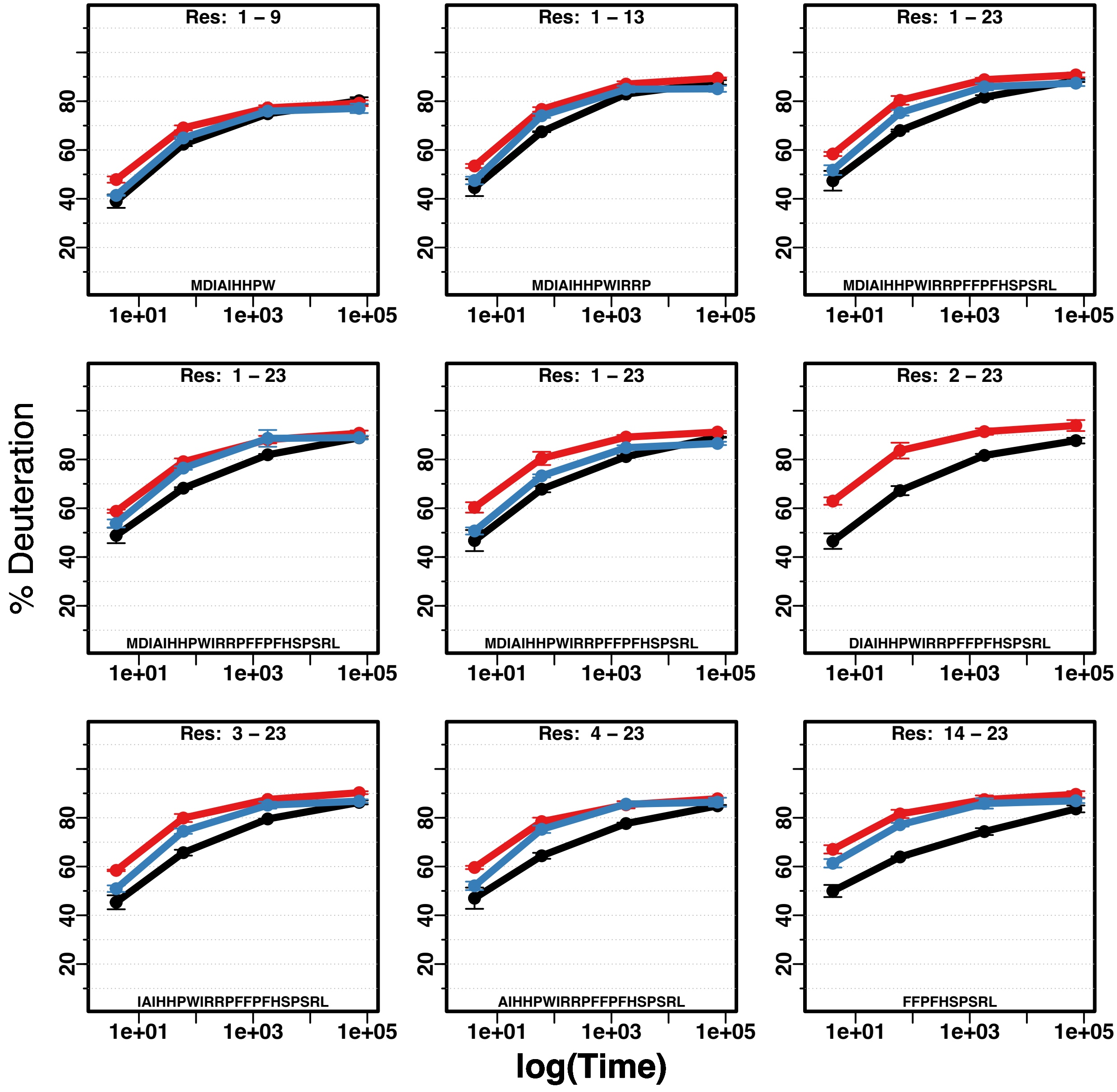

Res: 1 – 9
MDIAIHHPW
Res: 1 – 13
MDIAIHHPWIRRP
Res: 1 – 23
MDIAIHHPWIRRPFFFPFHSPSRL
Res: 1 – 23
MDIAIHHPWIRRPFFFPFHSPSRL
Res: 1 – 23
MDIAIHHPWIRRPFFFPFHSPSRL
Res: 2 – 23
DIAIHHPWIRRPFFFPFHSPSRL
Res: 3 – 23
IAIHHPWIRRPFFFPFHSPSRL
Res: 4 – 23
AIHHPWIRRPFFFPFHSPSRL
Res: 14 – 23
FFPFHSPSRL
% Deuteration
log(Time)
20
40
60
80
1e+01
1e+03
1e+05

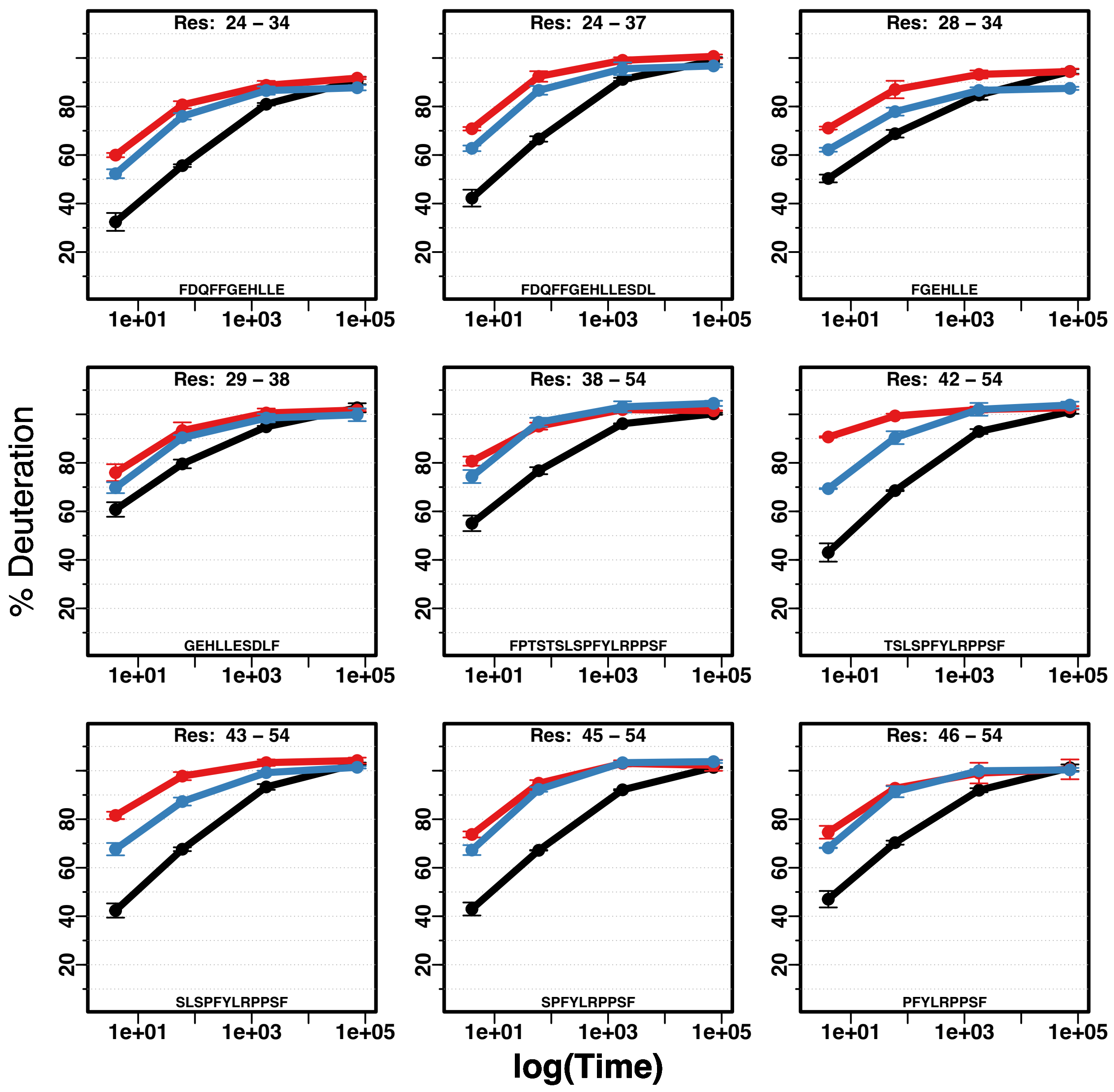

% Deuteration
log(Time)
Res: 24 – 34
FDQFFGEHLLE
Res: 24 – 37
FDQFFGEHLLESDL
Res: 28 – 34
FGEHLLE
Res: 29 – 38
GEHLLESDLF
Res: 38 – 54
FPTSTSLSPFYLRPPSF
Res: 42 – 54
TSLSPFYLRPPSF
Res: 43 – 54
SLSPFYLRPPSF
Res: 45 – 54
SPFYLRPPSF
Res: 46 – 54
PFYLRPPSF
20
40
60
80
1e+01
1e+03
1e+05

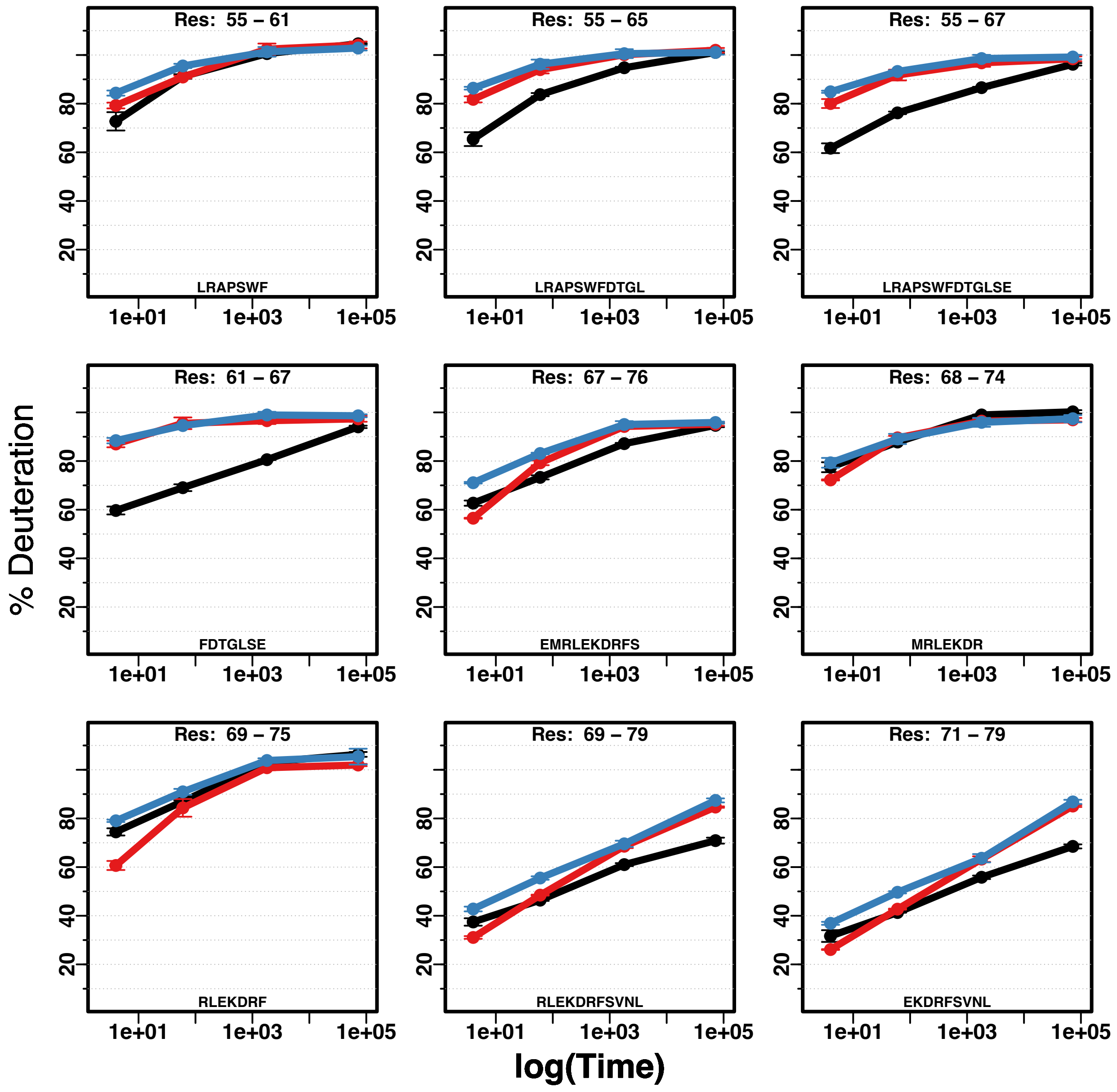

% Deuteration
log(Time)
Res: 55 – 61
LRAPSWF
Res: 55 – 65
LRAPSWFDTGL
Res: 55 – 67
LRAPSWFDTGLSE
Res: 61 – 67
FDTGLSE
Res: 67 – 76
EMRLEKDRFS
Res: 68 – 74
MRLEKDR
Res: 69 – 75
RLEKDRF
Res: 69 – 79
RLEKDRFSVNL
Res: 71 – 79
EKDRFSVNL
20
40
60
80
1e+01
1e+03
1e+05

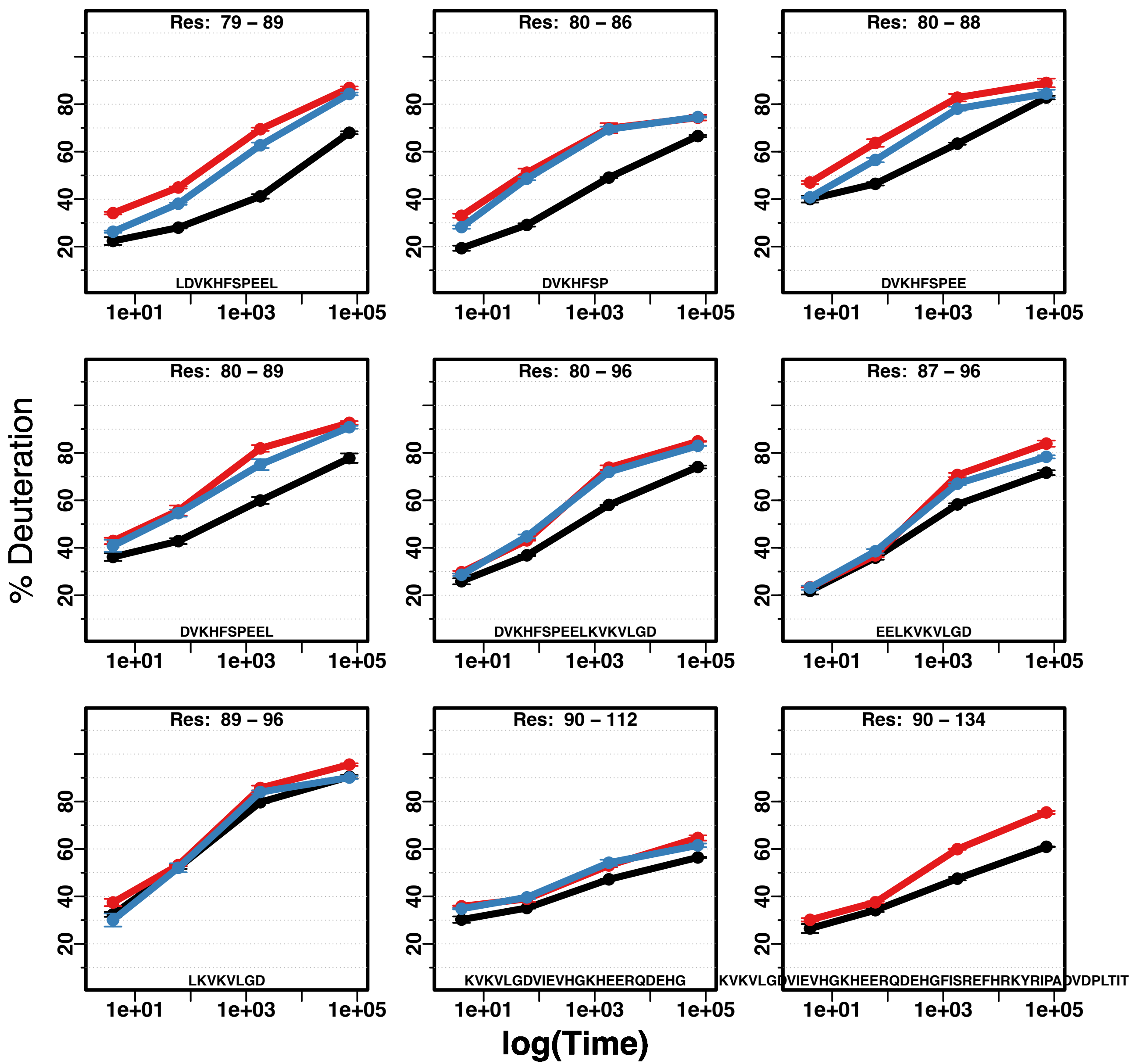
Res: 79 – 89
LDVKHFSPEEL
Res: 80 – 86
DVKHFSP
Res: 80 – 88
DVKHFSPEE
Res: 80 – 89
DVKHFSPEEL
Res: 80 – 96
DVKHFSPEELKVKVLGD
Res: 87 – 96
EELKVKVLGD
Res: 89 – 96
LKVKVLGD
Res: 90 – 112
KVKVLGDVIEVHGKHEERQDEHG
Res: 90 – 134
KVKVLGDVIEVHGKHEERQDEHGFISREFHRKYRIPADVDPLTIT
% Deuteration
log(Time)
20
40
60
80
1e+01
1e+03
1e+05

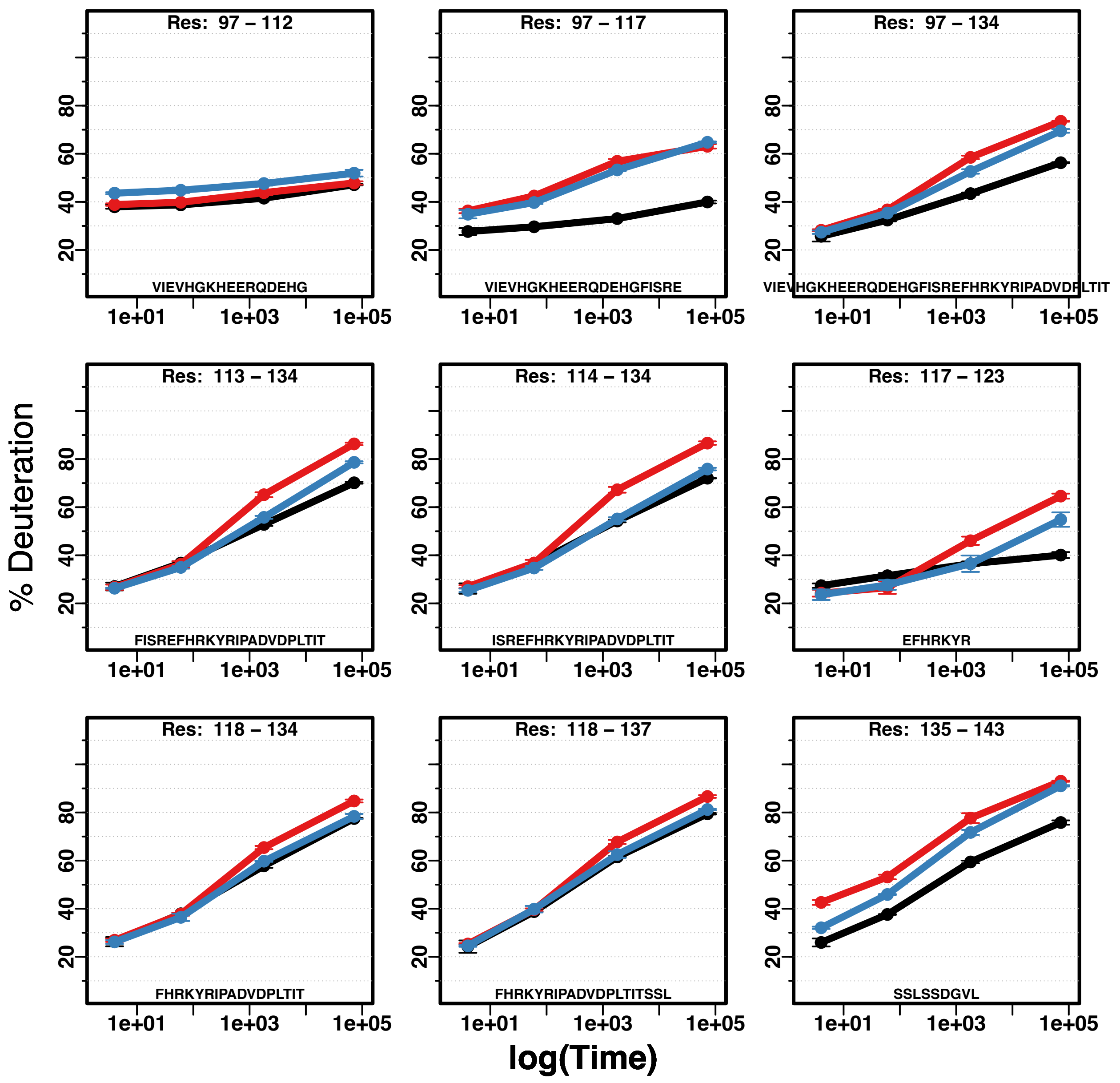
Res: 97 – 112
VIEVHGKHEERQDEHG
Res: 97 – 117
VIEVHGKHEERQDEHGFISRE
Res: 97 – 134
VIEVHGKHEERQDEHGFISREFHRKYRIPADVDPLTIT
Res: 113 – 134
FISREFHRKYRIPADVDPLTIT
Res: 114 – 134
ISREFHRKYRIPADVDPLTIT
Res: 117 – 123
EFHRKYR
Res: 118 – 134
FHRKYRIPADVDPLTIT
Res: 118 – 137
FHRKYRIPADVDPLTITSSL
Res: 135 – 143
SSLSSDGVL
% Deuteration
log(Time)
20
40
60
80
1e+01
1e+03
1e+05

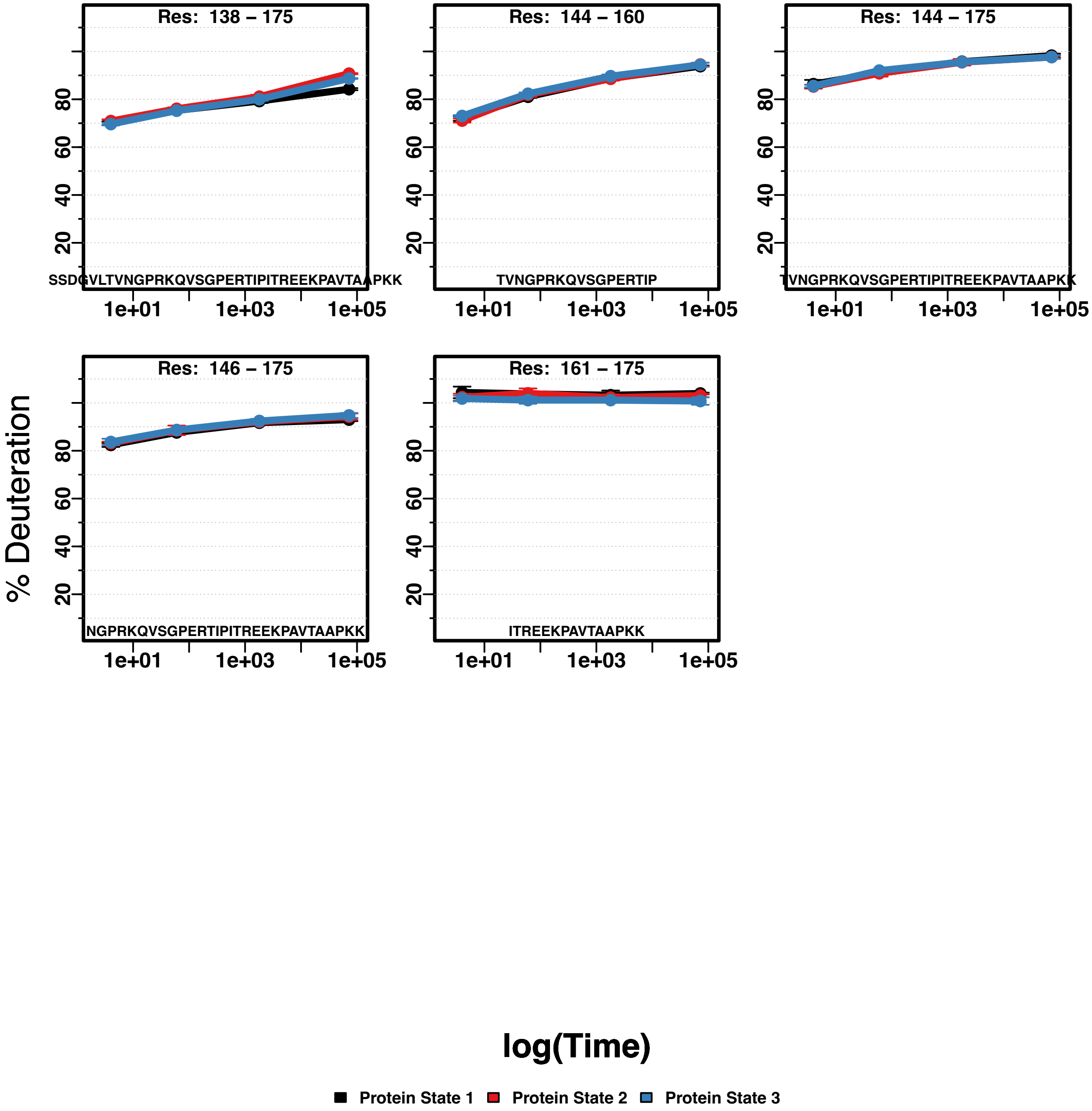
Res: 138 – 175
SSDGVLTVNGPRKQVSGPERTIPITREEKPAVTAAPKK
Res: 144 – 160
TVNGPRKQVSGPERTIP
Res: 144 – 175
TVNGPRKQVSGPERTIPITREEKPAVTAAPKK
Res: 146 – 175
NGPRKQVSGPERTIPITREEKPAVTAAPKK
Res: 161 – 175
ITREEKPAVTAAPKK
% Deuteration
20
40
60
80
1e+01
1e+03
1e+05
log(Time)
Protein State 1
Protein State 2
Protein State 3

Fig. S2. Uptake deuterium plots for all peptides of WT-, R120G-, and D109H-HSPB5. Timepoints are 4 sec, 60 sec, 1800 sec, 72000 sec. Statistics for HDX data are provided in Supplemental Table 1. Curves in black, red, and blue are for WT-, D109H, and R120G, respectively.

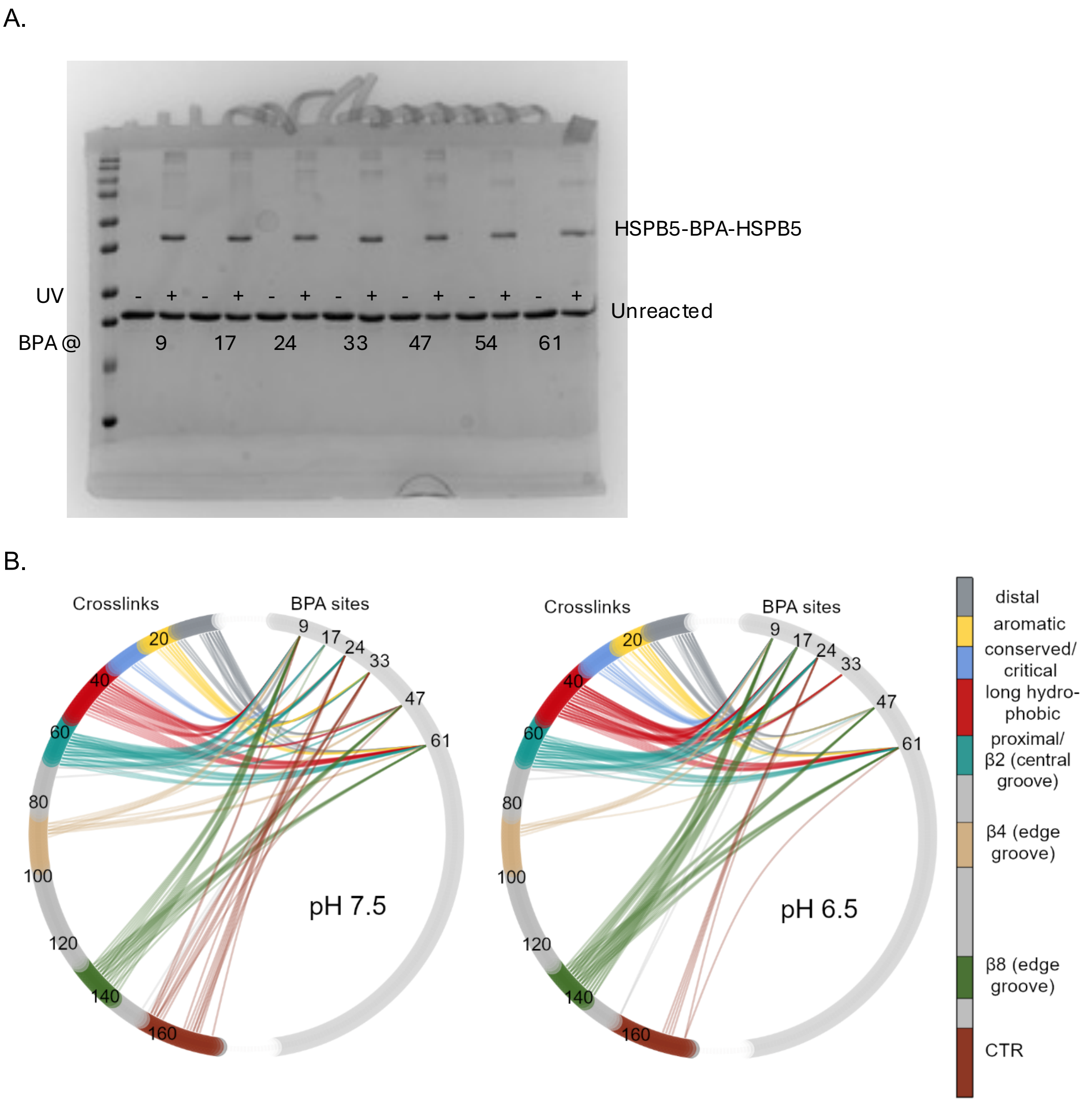


Fig. S3. Photo-crosslinking in HSPB5 oligomers. A) Example of products formed by UV irradiation of BPA-containing HSPB5 oligomers at pH 6.5. Bands marked "HSPB5-BPA-HSPB5" were subjected to in-gel proteolytic digestion and analyzed by MS. B) Circle Plot representation showing all identified crosslinks from six BPA variants. The BPA sites are on the right half of a circle; the identified crosslinks are on the left side of a circle. The colors refer to sub-regions and structural elements, as labeled.

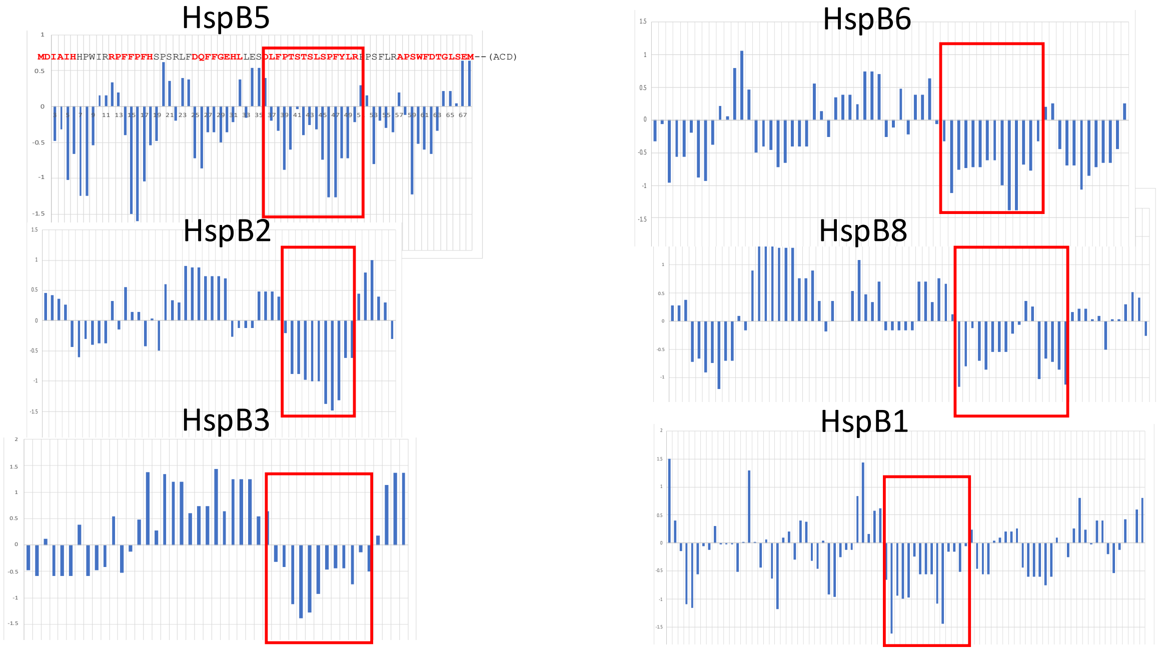


Fig. S4. Hydrophobicity plots of the NTRs of human sHSPs. The NTR sequences from six human sHSPs were analyzed using Hopp & Woods hydrophobicity scale with a window size of 5 residues. The sequence above HSPB5 is colored to show residues identified in crosslinking experiments. Red boxes identify the Long Hydrophobic region of HSPB5 and similar long regions of hydrophobicity near the proximal end of the other sHSPs.

**Table S1. Statistics for HDX data.**

| Protein | # reps | # peptides | Peptide Coverage (%) | <Peptide Length> | <Red> | St. dev. | CI | <Back HX> | Back HX Range |
|---|---|---|---|---|---|---|---|---|---|
| WT-HSPB5 | 3 | 50 | 100 | 14.84 | 4.93 | 0.074 | 0.293 | 71.7 | 36.52-87.98 |
| R120G-HSPB5 | 3 | 50 | 100 | 14.84 | 4.93 | 0.074 | 0.257 | 70.85 | 36.67-87.84 |
| D109H-HSPB5 | 3 | 48 | 100 | 14.1 | 4.49 | 0.071 | 0.239 | 70.32 | 36.18-87.61 |

Table S2. Comparison of Crosslinks Identified by BPA variant under non-activating (pH 7.5) and activating (pH 6.5) conditions.

| BPA site (Region) | Jensen-Shannon Similarity Score | Total Crosslink PSM (pH 7.5) | Total Crosslink PSM (pH 6.5) | % total by zone (pH 7.5) | % total by zone (pH 6.5) | # zones (pH 7.5) | # zones (pH 6.5) |
|---|---|---|---|---|---|---|---|
| 9 (Distal) | 0.76 | 215 | 195 | Long H 15<br>Prox/β2 35<br>Edge 45 | Long H 20<br>Prox/β2 24<br>Edge 50 | 3 | 3 |
| 17 (Aromatic) | 0.47 | 45 | 169 | Prox/β2 93 | Distal 25<br>Prox/β2 55<br>Edge 15 | 1 | 3 |
| 24 (Conserved) | 0.56 | 66 | 136 | Distal 18<br>Prox/β2 55<br>CTR 17 | Distal 28<br>Aromatic 29<br>Long H 14<br>Prox/β2 26 | 3 | 4 |
| 33 (Critical) | 0.51 | 62 | 42 | Distal 50<br>Aromatic 39 | Distal 17<br>Aromatic 69<br>Long H 10 | 2 | 3 |
| 47 (Long Hydrophobic) | 0.63 | 61 | 34 | Distal 51<br>Edge 39 | Distal 59<br>Prox/β2 15<br>Edge 24 | 2 | 3 |
| 61 (Proximal/β2) | 0.74 | 281 | 179 | Distal 56<br>Aromatic 32 | Distal 46<br>Aromatic 32<br>Long H 16 | 2 | 3 |

1.
